# Astex-Faro: improving detection of ligand binding in cryo-EM maps through statistical baseline analysis

**DOI:** 10.64898/2026.11.03.763890

**Authors:** Federico P. de Isidro-Gómez, Carlos Oscar S. Sorzano, José María Carazo, Michael Saur

## Abstract

Cryogenic electron microscopy (cryo-EM) is emerging as a tool for large-scale ligand screening and fragment-based drug discovery, generating extensive collections of reconstructions. At the same time, the systematic inspection of local density variations remains challenging, particularly when identifying weak or partially occupied features. Here, we present Astex-Faro, a statistical framework that defines an expected specimen state from a reference ensemble and identifies significant structural deviations in a target map through comparison against this reference. Furthermore, the method allows the enhancement of weak or poorly resolved features, enabling the recovery of signals that would otherwise be difficult to interpret. Fourier Shell Coherence (FSCoh) is introduced as a metric that quantifies consistency across reconstruction ensembles. We show that Astex-Faro reliably detects subtle structural features and improves their interpretability across datasets spanning a broad range of experimental challenges, from highly homogeneous specimens to membrane proteins, low-occupancy ligands and fragment-screening experiments. Together, these developments provide a practical framework for cryo-EM screening, increasing confidence in hit identification and reducing the risk of missed binding events.

## 1. Introduction

Cryogenic electron microscopy (cryo-EM) has become a useful tool in the design of new medicines and is increasingly complementing the use of X-ray crystallography for structure-based drug discovery (SBDD) [1] [2] [3]. A particularly challenging use case in this field is its application to Fragment-based Drug Discovery (FBDD), where many hundreds of datasets are analysed to determine whether small molecular fragments (typically with molecular weight < 300 Da [4]) have bound to a protein of interest (the target). Due to the small size of fragments, interactions between fragment and target are generally weak, with binding affinities (K_d_) typically in the high micromolar to low millimolar regime, putting their detection by both biophysical and structural methods on the edge of what is technically feasible. In addition to their intrinsically small size, which can make identification challenging, the weak interaction between fragment and target often also leads to partial occupancy, with not all copies of the target molecule in the experimental sample having a fragment bound. Since both the traditionally used tool for FBDD, X-ray crystallography, and single particle cryo-EM are methods which analyse an average of many protein molecules, this partial occupancy weakens the ligand features in the final structure and further complicates the successful identification of binding events.

To increase the success rate of identifying such weak and partially populated binding events, a widely used tool in x-ray crystallography FBDD campaigns is the Pan-Dataset Density Analysis (PanDDA) [5]. PanDDA analyses a pool of apo data, i.e. structures which do not have any ligand bound, to determine which features are statistically present in the apo state. It then compares a new structure against this reference and highlights regions that differ significantly from the apo pool. In case of partial occupancy, PanDDA also weights statistically different features so that ligand features become clearer and easier to interpret. An equivalent application of this method for cryo-EM has not yet been released. Other approaches aim to estimate and normalize local contrast variations [6] [7], assess the fit of a ligand model to the density [8], or propose ligand candidates for a previously identified density [9], but do not provide pan-dataset identification of outlier features.

Here we present Astex-Faro, a new method which identifies regions within cryo-EM reconstructions that deviate from a reference set of maps. This method highlights those regions and determines a weighting factor used to improve the features that are statistically different from the reference pool. We also introduce Fourier Shell Coherence (FSCoh) as part of Astex-Faro, a metric necessary to analyse pools of maps and specifically designed to evaluate multi-map information consistency. The purpose of Astex-Faro is to reduce the number of binding events that are missed during intensive cryo-EM screening campaigns, provide confidence to identify such events as genuine and, hence, to increase the screening output. We evaluated the method on three targets, *β*-Galactosidase (Bgal) [1], transient receptor potential mucolipin 1 (TRPML1) [10], and Pyruvate Kinase M2 (PKM2) [1] (see Supplementary Tables 1 and 2).

## 2. Results

### 2.1. FSCoh measures multi-map information consistency

Astex-Faro relies on a collection of maps that provide the reference population against which individual maps are evaluated. Inherent variability in cryo-EM experiments can lead to differences in the final maps produced for a dataset, even when using the same sample type and imaging conditions. Hence, it is important to determine up to which resolution the members of the map pool remain mutually consistent, such that the statistical analysis is based only on information consistently supported by the ensemble. In cryo-EM, the Fourier Shell Correlation (FSC) quantifies agreement between two maps by providing a global measure of the highest resolution at which they share information, given a cut-off threshold [11]. Because FSC is inherently a pairwise measure, it cannot directly assess agreement across multiple maps.

We propose Fourier Shell Coherence, a new measure of the spectral agreement between any number of maps (see Section 4.2 for the formal definition). Coherence provides a measure of how consistently information is shared across multiple signals, treating the ensemble as a single entity. FSCoh generates a spectral coherence profile and quantifies coherence at each spatial frequency across the map pool.

Figure 1a, Figure 2a, and Figure 3a show the FSCoh curves calculated from pools of apo Bgal, TRPML1, and PKM2 maps, respectively, along with the FSC curves derived from the half-maps of each reconstruction in the corresponding pool. For the Bgal pool, individual FSC resolutions range from 2.7 to 3.7 Å, while FSCoh reports a pool-wide consistency of 3.3 Å, consistent with the dataset’s overall resolution range. Similar trends are observed for TRPML1 and PKM2. The TRPML1 pool of 10 maps shows FSC resolutions between 2.5 and 2.9 Å, while FSCoh reports a pool-wide consistency of 2.8 Å. The PKM2 pool of 10 maps shows individual FSC resolutions ranging from 3.1 to 3.8 Å, while FSCoh reports a pool-wide consistency of 3.7 Å. This last specimen shows the strongest signal variability, as reflected by the FSCoh.

**Figure 1.**
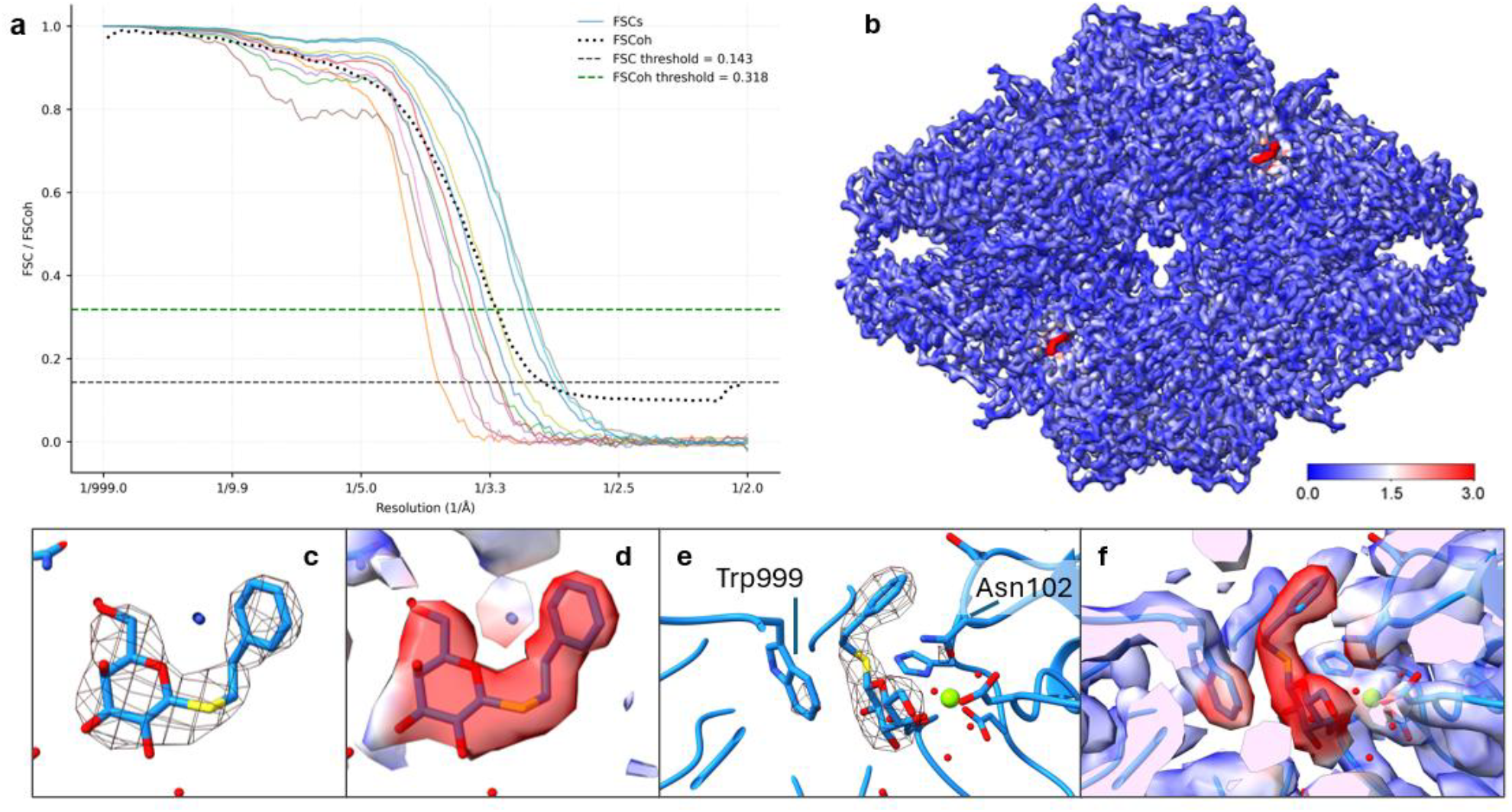
Analysis Bgal ligand binding. (a) Individual FSCs and FSCoh curves used to assess map quality and define a common resolution threshold for the reference pool. Individual FSC curves indicate mapspecific resolution estimates between 2.7 and 3.7 Å, while the FSCoh curve defines a pool-wide cutoff at 3.3 Å. (b) Statistical map of Bgal in complex with compound B1. The z-score values used for colouring are indicated in the colour bar. Note the red colouring of the ligand feature, highlighting significant deviation from the reference apo pool. (c,e) Z-score map focused on the binding site at a threshold of 3 (d,f) Statistical map focused on the binding site; colouring same as in (b). Note that both the z-score map and the statistical map show statistically significant changes in some of the binding pocket side chains (Trp999 and Asn102) as well, consistent with binding-associated side-chain reordering.

**Figure 2.**
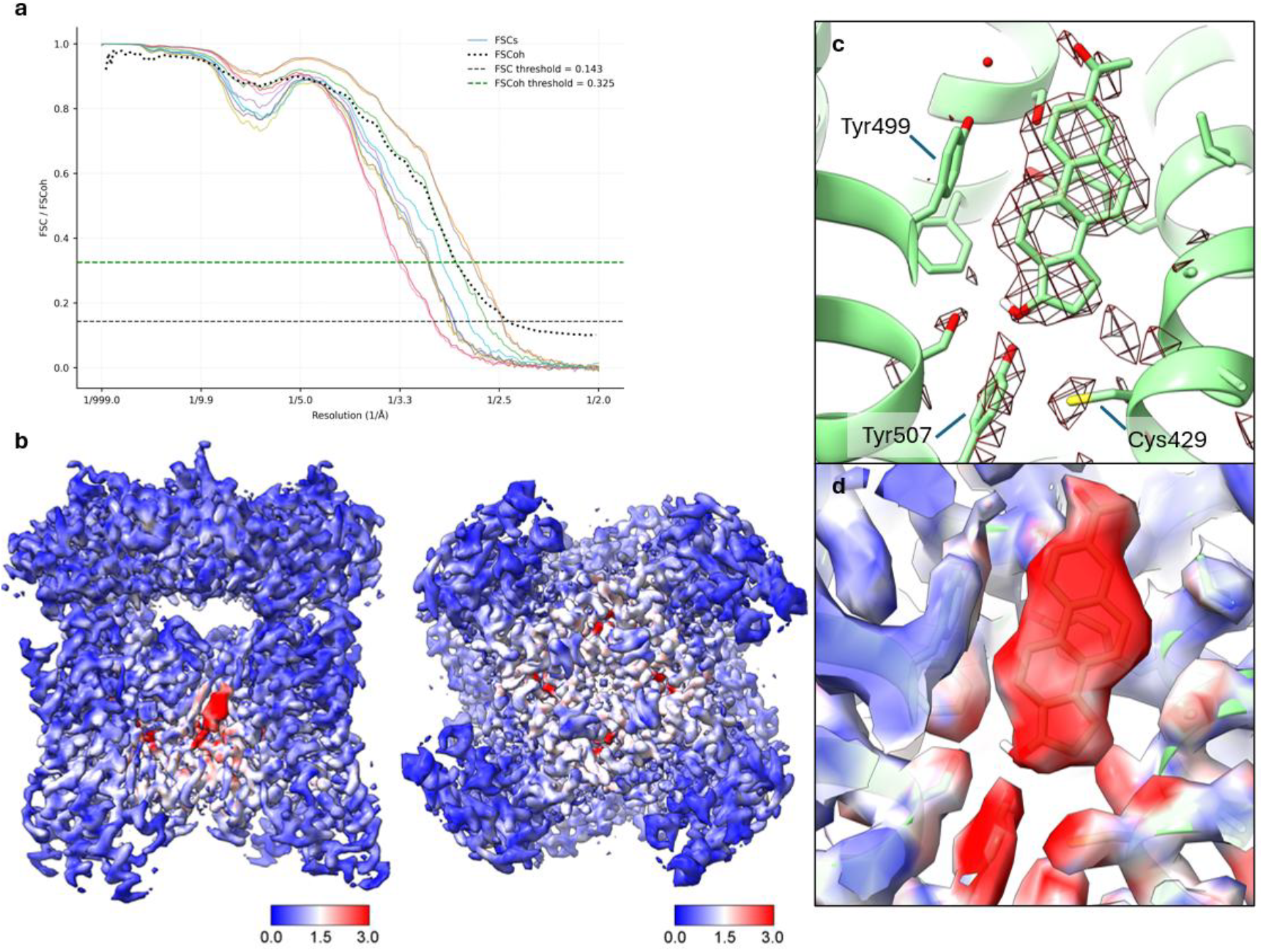
Analysis of TRPML1 ligand binding to compound T1. (a) Individual FSCs and FSCoh curves used to assess map quality and define a common resolution threshold for the reference pool. Individual FSC curves indicate map-specific resolution estimates between 2.5 and 2.9 Å, while the FSCoh curve defines a pool-wide cutoff at 2.8 Å. (b) Statistical map of TRPML1 in complex with compound T1 in side and top views. The z-score values used for colouring are indicated in the colour bar. Note the red colouring of the ligand located in the transmembrane region, highlighting significant deviation from the reference apo pool. (c) z-score map focused on the binding site at a threshold of 3 (d) Statistical map focused on the binding site; colouring same as in (b). Note that both z-score map and statistical map show statistically significant changes in some of the binding pocket side chains (Tyr 507 and Cys429) as well, consistent with binding-associated side-chain reordering.

**Figure 3.**
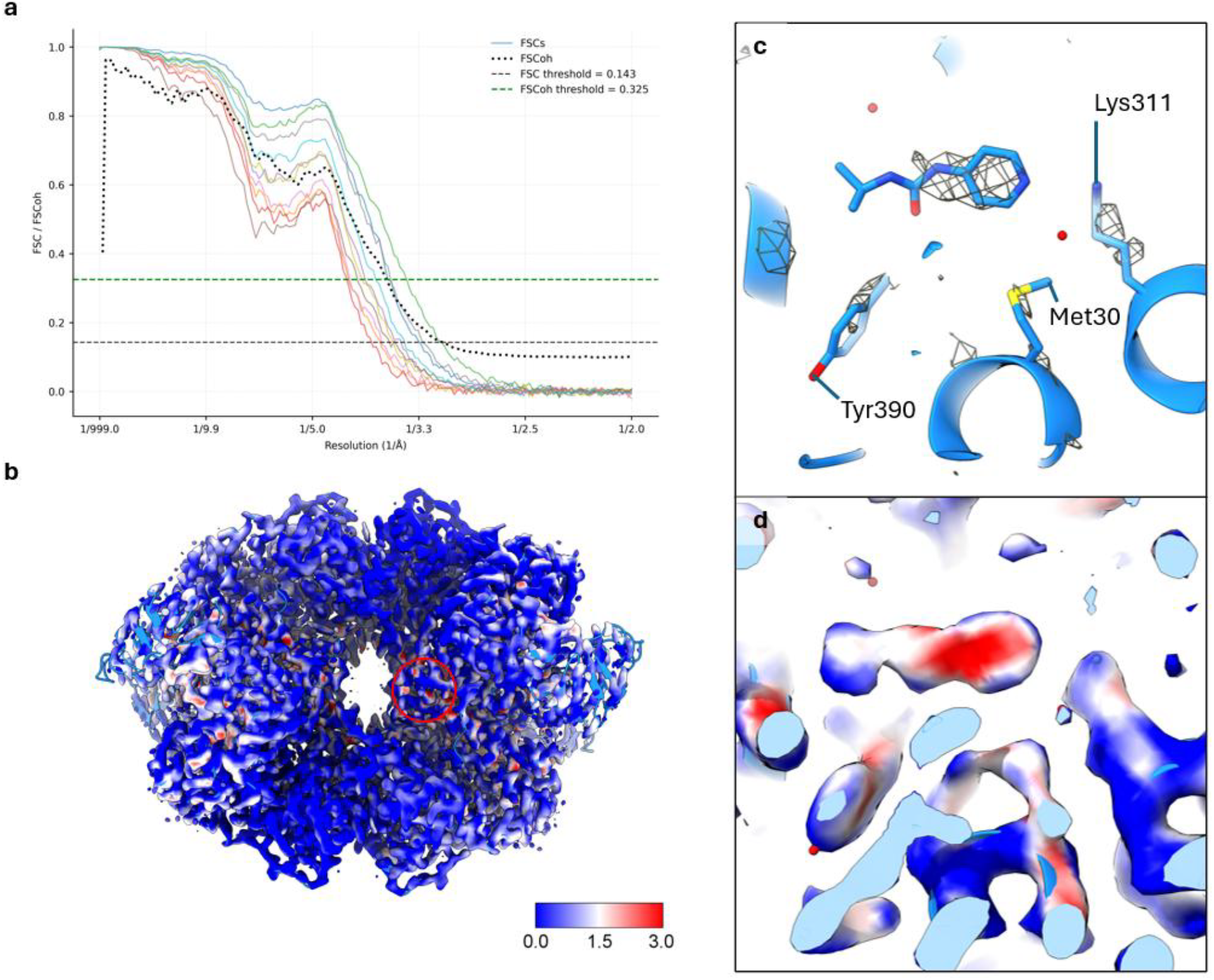
Analysis of PKM2 ligand binding. (a) Individual FSCs and FSCoh curves used to assess map quality and define a common resolution threshold for the reference pool. Individual FSC curves indicate map-specific resolution estimates between 3.1 and 3.8 Å, while the FSCoh curve defines a pool-wide cutoff at 3.7 Å. (b) Statistical map of PKM2 in complex with compound P1. The z-score values used for colouring are indicated in the colour bar. Note the red colouring of the ligand, highlighting significant deviation from the reference apo pool. (c) z-score map focused on the binding site at a threshold of 3. (d) Statistical map focused on the binding site; colouring same as in (b).

### 2.2. FSCoh relates to widely accepted FSC signal threshold

The agreement between FSCoh and conventional FSC measurements is not merely empirical. Both metrics can be expressed in terms of signal-to-noise ratio (SNR), becoming directly comparable within a common mathematical framework. Thus, we establish a direct mathematical relationship between the metrics, linking pairwise and ensemble-based assessments of signal consistency (see Section 4.2).

Both FSC and FSCoh require a threshold to determine map consistency. For FSC, the 0.143 criterion [11] has become the *de facto* standard in cryo-EM. Based on the relationship between FSC and signal-to-noise ratio (SNR). Through the direct mathematical correspondence between FSC and FSCoh, any FSC threshold can be translated to its equivalent FSCoh criterion, which depends solely on the number of maps in the ensemble (see Section 4.3 and FSCoh Supplementary Sections 2 to 4). As a result, FSCoh inherits the consistency criterion defined in the FSC framework while extending its applicability from pairwise to multi-map analyses. This direct connection with the FSC formalism provides a principled foundation for FSCoh.

Under this framework, FSCoh values estimated directly from reconstruction ensembles are expected to agree with those predicted from the corresponding individual FSCs. To validate this relationship, we conducted two complementary analyses. First, we performed controlled simulations on data with known signal and noise properties. Second, we analysed an ensemble of 10 independent reconstructions obtained from a single microscopy acquisition, providing a highly consistent experimental dataset. In both settings, the observed FSCoh values closely matched the predictions derived from the FSC framework. These results provide empirical support for the theoretical relationship between the two metrics and for the proposed thresholding scheme (see FSCoh Supplementary Section 6).

A direct consequence of the FSCoh formalism is that differences between the values calculated from the individual FSCs and the empirically measured FSCoh arise from disparities in signal content across the reconstruction ensemble. Such deviations indicate that reconstructions differ not only due to noise but also due to variations in the underlying signal, reflecting conformational heterogeneity within the pool or alignment and contrast inconsistencies between maps (see Supplementary Figure 6).

### 2.3. Astex-Faro highlights ligand binding events

Astex-Faro characterises a pool of reconstructions and establishes a statistical model of the reference ensemble. Comparison of a target map against the reference produces a z-score map, where each voxel quantifies the deviation of the observed density from the expected reference distribution. This is calculated as the difference from the reference mean, normalized by the combined local standard deviation and a global variability term derived from the median absolute deviation (MAD). Regions with z-scores exceeding the significance threshold (z-score > 3) are classified as unexpected densities in the target map. When the z-score map is used to colour the target reconstruction, it provides a statistical map that highlights regions deviating from the reference distribution (see Section 4.4).

Figure 1 shows the results of statistically analysing a pool of apo Bgal maps against a map of Bgal in complex with compound B1. The experiment used in-house data comprising 11 apo maps as a reference pool and one holo map as a target map, which was evaluated against the reference pool. FSC curves span a range between 2.7 and 3.7 Å for the individual reconstructions, consistent with the 3.3 Å overall resolution for the map pool reported by the FSCoh (Figure 1a). We then filtered all maps to the FSCoh threshold, ensuring downstream analyses used a common, reliable resolution band. The statistical map (Figure 1b) shows significant deviations from the reference pool in both the ligand and its binding pocket. In particular, the ligand density is clearly distinguishable from the reference pool (Figure 1 d,f), and its region is accurately delineated by the z-score map (Figure 1 c,e). Notably, the analysis also identifies the displacement in two side chains (Trp999 and Asn102) within the pocket that are associated with ligand interaction (Figure 1 e & f) [1].

We also evaluated Astex-Faro on the membrane protein TRPML1. This system is more challenging because of its intrinsic structural heterogeneity and the surrounding lipid bilayer, with the binding pocket located within the transmembrane region. Compared to the Bgal/compound B1 complex, ligand binding features in TRPML1 show lower contrast, and the density corresponding to compound T2 is less continuous (see Section 2.4). In addition, the TRPML1 binding site in the apo state shows noticeable residual density, whereas Bgal provides a comparatively cleaner binding region (see Supplementary Figure 5). Figure 2 shows the results of statistically analysing a previously published dataset of TRPML1 in complex with compound T1 [10]. The complex was compared to a pool of 10 apo maps with a resolution range of 2.5-2.9 Å and a common FSCoh resolution of 2.8Å. The resulting statistical map shows significant variability at both the ligand site and across the binding pocket (Figure 2b). Subtle variability can also be observed throughout the transmembrane region, reflecting ligand-induced heterogeneity, with more pronounced variability visible in the pore region. Despite the high variability and increased complexity imposed by the detergent micelle, Astex-Faro identifies subtle but meaningful structural features in the transmembrane region as statistically significant, including ligand binding and ligand-induced side chain rearrangements (Figure 2 c & d).

To determine if Astex-Faro is sensitive enough to detect the binding of fragment-sized molecules we analysed a previously published dataset of PKM2 in complex with compound P1 [1]. The apo pool of maps comprising 10 structures showed a per-map FSC resolution range of 3.1-3.8 Å and FSCoh resolution of 3.7 Å. The statistical map shows obvious variability in the binding site and surrounding regions, with compound P1 clearly marked as a statistically relevant deviation from the apo pool of maps (Figure 3). During a fragment screen, this protein-wide highlighting of statistical deviation from the apo state could provide helpful markers to aid the identification of binding events where the quick localisation of a small additional map feature in the shape and size of a fragment might prove challenging. Despite the ligand feature clearly being marked as significantly different from the apo state, Figure 3d also shows that parts of the feature are marked as less significant changes from apo than others. This is explained by residual features present in the binding site of the apo map pool which partially overlay the ligand in the target map. Accordingly, regions of compound P1 that coincide with these residual apo features display a reduced statistical significance, whereas the remaining regions are strongly significant. These results demonstrate Astex-Faro sensitivity to ligand-induced changes in sites of interest despite pre-existing apo state features.

Finally, to demonstrate the robustness of the method against the intrinsic variability and presence of noise in cryo-EM reconstructions, an apo reconstruction which was not present in the reference pool was analysed against the apo reference ensemble. No statistically significant changes were detected, indicating that the reference ensemble accurately captures the apo state (see Supplementary Figure 8).

### 2.4. Astex-Faro can improve ligand features for partially occupied binding sites

We have shown that Astex-Faro highlights features in a target map that are statistically different from a reference pool of maps and that this analysis can be used to highlight ligand binding events (Section 2.3). Weak ligands can show partial binding to proteins, leading to weak ligand features in the final reconstruction of the protein-ligand complex. We hypothesised that those ligand features can be improved by estimating and applying a weighting factor that removes apo information from the map (see Section 4.5 for details). To test this hypothesis, we combined predefined proportions of particles from Bgal apo and liganded datasets *in silico* to generate maps corresponding to distinct partial occupancy conditions (Figure 4, Supplementary Figure 7).

**Figure 4.**
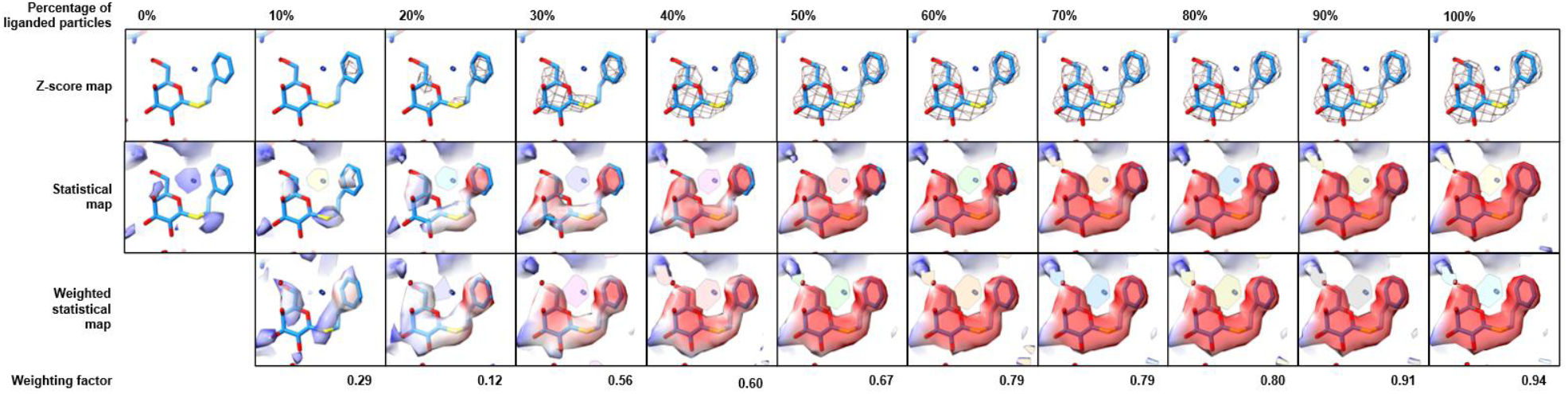
Effect of partial occupancy weighting on ligand feature quality Bgal in complex with compound B1. In silico mixing of apo and liganded particles, ranging from 0% to 100% liganded data, in 10% steps. Each column represents one dataset with a specific mixing ratio. First row: amount of liganded particles in each dataset. Second row: z-score map for each dataset, focused on the ligand. Third row: corresponding statistical map for each dataset. Fourth row: statistical maps weighted by the weighting factor shown in the fifth row. The weighted map for the 0% bound-particle condition is not shown, as no statistically significant regions are detected and no weighting factor can be calculated. An equivalent figure showing a side view of the ligand is provided in Supplementary Figure 7.

Eleven mixed datasets were generated, ranging from 0% to 100% liganded particles in steps of 10%, while keeping the total number of particles constant across datasets (Supplementary Table 3). To avoid alignment-driven artifacts from previous reconstructions, particle orientations were estimated by subjecting each mixed dataset to 3-D refinement. Figure 4 shows that as the fraction of bound particles decreases, ligand features progressively degrade and statistically relevant differences to the apo pool of maps decrease. Applying the calculated weighting factor to the map effectively recovers the weakened ligand signal. This is particularly evident at low occupancy levels of 20% and 30%, where ligand connectivity is recovered.

The performance of the method was further evaluated by analysing a dataset of TRPML1 in complex with compound T2 [10]. This ligand exhibits weaker ligand features than Bgal/B1, TRPML1/T1 and PKM2/P1, with features disappearing more quickly than surrounding side chain features when increasing the voxel intensity threshold at which the structure is shown (Figure 5 a-c). This suggests partial occupancy of compound T2. Accordingly, the weighting factor calculated by Astex-Faro is 0.46, and rescaling the target map reveals stronger ligand features, with ligand connectivity being established at higher thresholds (Figure 5d-f). Furthermore, as in PKM2/P1 (Section 2.3), not all ligand features in the map exhibit the same level of statistical significance (Figure 5 c). Relative to Bgal/ B1, compound T2 occupies a slightly different position within the TRPML1 antagonist-binding site, partially overlapping residual features present in the apo map pool (Supplementary Figure 5). Despite these residual apo-state features, Astex-Faro correctly highlights the statistically relevant changes in the binding pocket introduced by compound T2.

**Figure 5.**
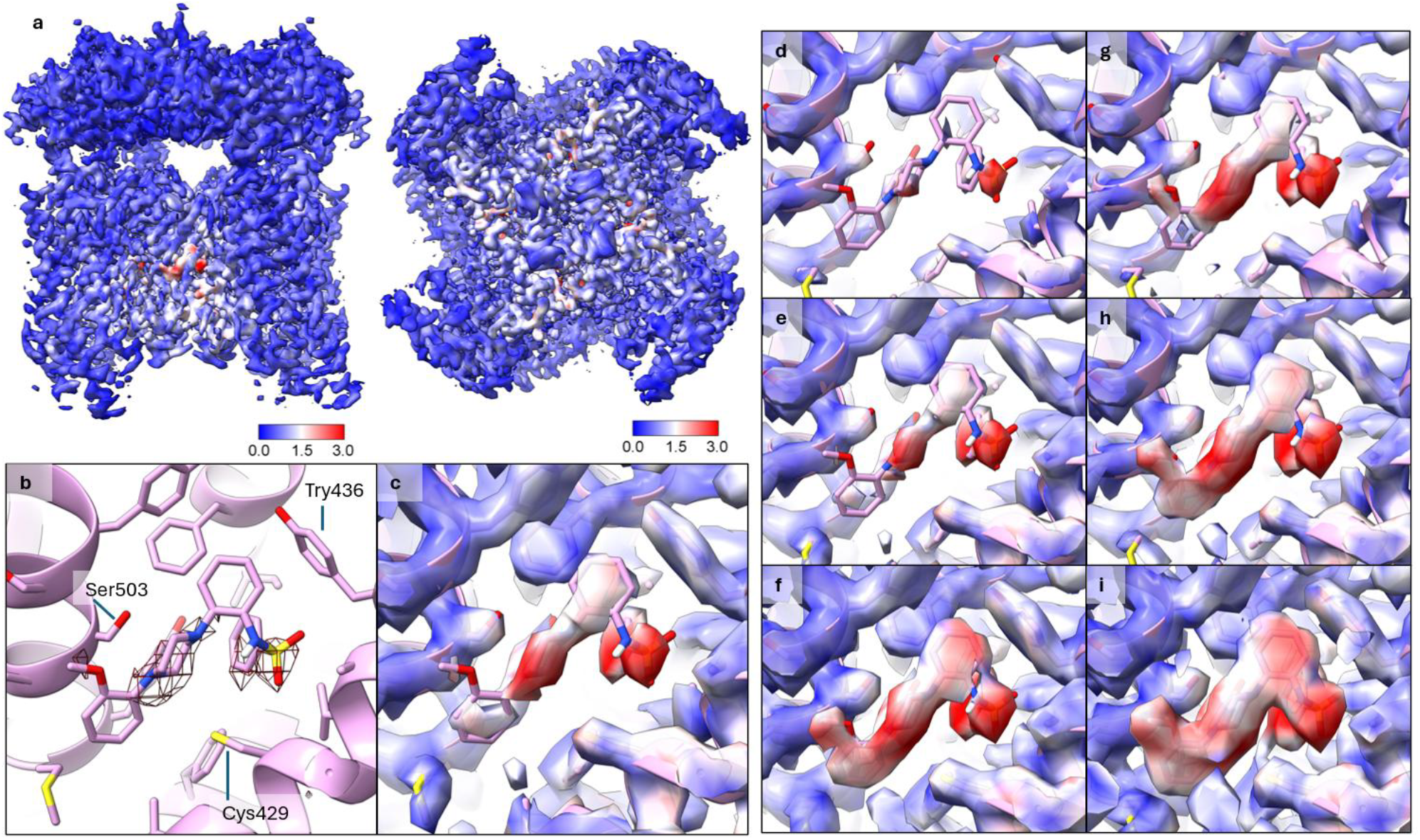
Analysis of TRPML1 ligand binding to compound T2. (a) Statistical map of TRPML1 in complex with compound T2 in side and top views. The z-score values used for colouring are indicated in the colour bar. (b) z-score map focused on the binding site at a threshold of 3. (c) Statistical map focused on the binding site; colouring is the same as in b). (b–f) original map at three different voxel intensity thresholds (2.5, 1.9, and 1.3), illustrating how the ligand density disappears at higher thresholds than the surrounding protein side chains. (g–i) weighted map at thresholds (1.6, 1.1, and 0.7) matching protein intensity as in d-f, illustrating how map weighting makes ligand density comparable to the contrast on the rest of the protein.

To test the limit of compound T2 detectability and ligand feature recovery by map weighting, we performed an *in silico* particle mixing experiment analogous to the one for Bgal (Supplementary Table 3). Given that ligand feature of compound T2 already shows low contrast in the original dataset, contrast of the ligand feature rapidly decreased with increasing ratios of apo particles (Figure 6). As expected, this progressively compromised the detection of statistically significant regions. The method reached its detection limit when the fraction of apo particles reached 50%, at which point no statistically significant ligand region was identifiable and, consequently, no weighted map could be calculated. However, whenever ligand features remained detectable, map weighting effectively enhanced ligand signal, restored connectivity, and balanced protein and ligand voxel intensities (Figure 6, bottom row).

**Figure 6.**
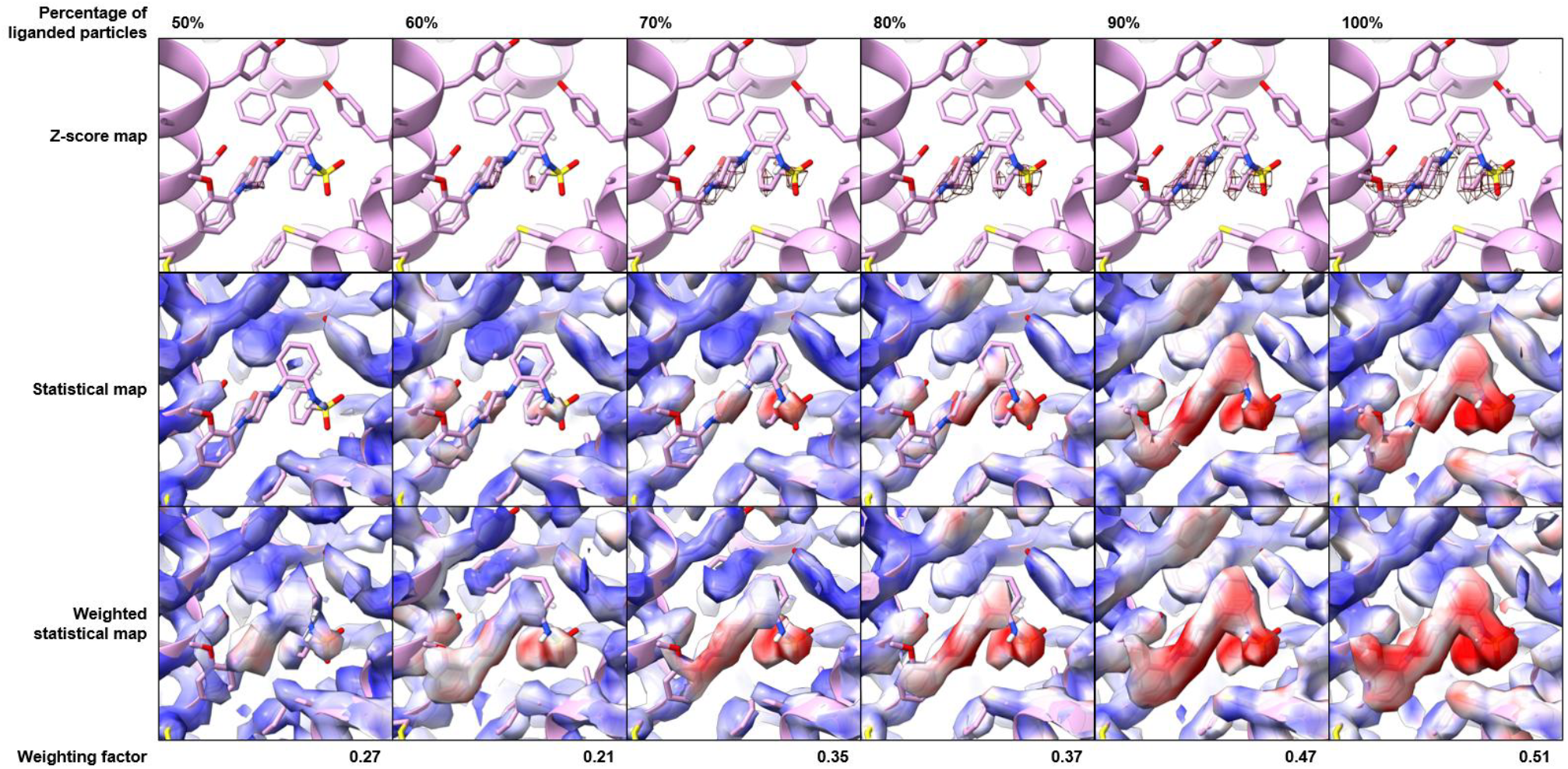
Effect of partial occupancy weighting on ligand feature quality in TRPML1/compound T2. In silico mixing of apo and liganded particles, ranging from 50% to 100% liganded data, in 10% steps. Each column represents one dataset with a specific mixing ratio. First row: amount of liganded particles in each dataset. Second row: z-score map for each dataset, focused on the ligand. Third row: corresponding statistical map. Fourth row: statistical maps weighted by the weighting factor shown in the fifth row.

From the conducted experiments, weighting-factor estimates displayed a more unstable behaviour at lower occupancy levels. Because both the ligand signal and the extent of the statistically significant region change as occupancy decreases, their respective contributions to weighting-factor estimation are difficult to disentangle. To better understand the contribution of each factor, we performed an additional analysis in which weighting factors were estimated using the region detected at full occupancy while varying the fraction of liganded particles. Using a fixed region resulted in more stable weighting-factor estimates, suggesting that part of the observed variability arises from changes in the detected region rather than from the ligand signal alone (see Supplementary Material Section 2).

### 2.5. Benchmark against pre-existing methods

Occupy [6] aims to estimate and normalise local signal variations without performing statistical pandataset analyses. We compared Occupy’s weighting of the TRPML1/compound T2 dataset with the results obtained using Astex-Faro. Occupy does not identify statistically significant outlier regions. Therefore, for comparison purposes, the ligand region analysed by Occupy was defined using the outlier mask identified by Astex-Faro. Within this region, the mean Occupy value was 0.54, in close agreement with the weighting factor estimated by Astex-Faro (0.46). Occupy also provided effective global density normalisation across the reconstruction (see Supplementary Material Section 1a).

We also evaluated LocScale2 [7], a method for local sharpening and density modification that aims to improve map interpretability. The method was evaluated using both feature-enhancement map (FEM) and model-free modes. Similar to Occupy, the method improved contrast homogeneity throughout the reconstruction, particularly in regions displaying weaker protein density. In the TRPML1/compound T2 dataset, enhanced density was observed across the transmembrane domain, with particularly strong effects in membrane-associated regions when using the FEM mode (see Supplementary Material Section 1b).

Despite these improvements in overall map contrast and interpretability, neither method provides a statistical framework for evaluating whether recovered features are expected from a reference population. Consequently, neither approach directly quantifies the statistical significance of observed density variations nor specifically targets the recovery of ligand-associated features. In contrast, Astex-Faro derives weighting factors from a statistical comparison against a reference ensemble and focuses enhancement on regions identified as statistically significant deviations from the expected system state (see Supplementary Material Section 1c).

## 3. Discussion

We introduced Astex-Faro, a framework for identifying outlier densities in cryo-EM reconstructions relative to an expected system state defined by a reference pool of maps. The framework combines statistical map analysis with a weighting strategy for recovering weak features and introduces Fourier Shell Coherence (FSCoh), a metric for assessing consistency across ensembles of reconstructions. Together, these components provide an integrated approach for detecting and interpreting structural deviations from a reference state.

We demonstrated the applicability of Astex-Faro across a diverse set of cryo-EM datasets. Statistically significant ligand features were successfully identified in systems ranging from Bgal benchmark to the more challenging TRPML1 membrane protein datasets, including low-contrast ligand-binding events such as compound T2. The method also proved sufficiently sensitive to detect fragment binding in PKM2. In addition, both *in silico* and experimental partial-occupancy analyses showed that the proposed weighting strategy can recover ligand contrast and improve the interpretability of weak binding events while preserving the statistical framework used to detect them.

FSCoh arises naturally from the statistical framework presented in this work and provides a practical measure of consistency across a pool of reconstructions. By identifying the frequency range over which the reference ensemble remains self-consistent, FSCoh defines the bandwidth within which downstream statistical analyses can be performed reliably, removing frequency components that can lead to misleading interpretations. Through its direct mathematical relationship with the conventional FSC, FSCoh extends an established consistency criterion from pairwise comparisons to multi-map analyses.

The power of Astex-Faro relies on its ability to statistically characterize the state of a specimen from an ensemble of reconstructions. Once such a characterization is established, any target map can be systematically interrogated against the reference ensemble, revealing outlier features relative to the expected structural landscape. In the context of drug discovery, this increases confidence in ligand hit identification, improves interpretation of weak ligand density, and reduces the likelihood of failing to detect binding events.

A practical limitation of multi-map approaches in cryo-EM is the relatively small number of reconstructions typically available to describe a given biological state. Larger ensembles would provide a more robust statistical representation of the specimen. However, generating many independent reconstructions of the same state remains costly and experimentally demanding. Despite using reference pools containing only 10 to 11 maps, Astex-Faro produced robust and interpretable results across all tested systems, demonstrating that useful statistical characterisation can already be achieved with currently accessible cryo-EM datasets.

Together, these results demonstrate that Astex-Faro enables the systematic identification of statistically significant structural features in cryo-EM reconstructions and their recovery in cases of compromised contrast. Such features may reflect conformational changes, ligand-binding events, or other phenomena of relevance. By providing an objective statistical framework for their detection and interpretation, Astex-Faro increases confidence in ligand identification and reduces the risk of missed binding events during cryo-EM protein-ligand structure determination for both SBDD and FBDD campaigns.

## 4. Materials and Methods

### 4.1. Datasets acquisition and processing

All maps used in this study, including the apo reference maps and ligand-bound target maps, were generated in-house for the datasets presented: β-galactosidase, PKM2 [1] and TRPML1 [10]. Sample preparation, data collection and image processing have been described previously [1] [10]. Before applying the methods presented here, we resampled all maps to a common box size and pixel spacing (1 Å/px) and aligned them to a shared reference geometry using ChimeraX [12].

The *in silico* partial-occupancy experiments required additional processing, as occupancy mixtures cannot be reliably generated from the reconstructed maps alone. For each dataset, we reextracted particles from the final refinement iteration of the apo and ligand-bound reconstructions (at 1 Å/px). We chose apo datasets to closely match the resolution of their corresponding ligand-bound datasets. Mixed particle populations containing a fixed number of particles and varying proportions of ligand-bound and apo particles were then generated. To avoid alignment-driven artefacts arising from previous reconstructions, particle orientations were re-estimated for each mixed dataset by an independent 3D auto-refinement in RELION 4.0 (Bgal; experimental details in [13]) or cryoSPARC (TRPML1; experimental details in [14])

The Astex-Faro results presented in this work were obtained using default parameters and without providing region of interest masks (see Sections 4.4 and 4.5). The complete workflow, including FSCoh estimation, statistical characterisation of the reference ensemble and target map evaluation, required less than one minute on a single CPU core for the datasets analysed in this study.

### 4.2. Fourier Shell Coherence (FSCoh)

To evaluate the consistency of multiple cryo-EM reconstructions, the Fourier Shell Coherence (FSCoh) is defined as a spectral metric that informs the consistency of information across a set of reconstructions. The normalized coherence, *C*(*ω*) of *N* signals, 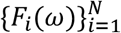, at frequency *ω* is defined as

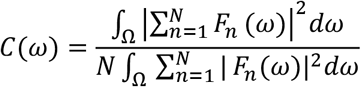

where *F*_*n*_(*ω*) denotes the Fourier component of the *n*-th map at frequency *ω* and 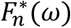 denotes the complex conjugate of a scalar Fourier component, averaged over shells Ω at frequency *ω*. In the discrete case, the integral can be replaced by the dot product of two row vectors containing the Fourier components:

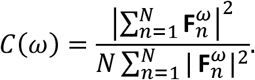

Under the assumption of additive noise, the FSCoh of *N* maps can be formulated as a perfect signal s corrupted by noise n_*i*_. Due to the relationship between the signal-to-noise ratio (SNR) and the Fourier Shell Correlation (FSC) [11], it is possible to establish a relationship between the FSCoh of the *N* maps and the individual FSC of each of them as

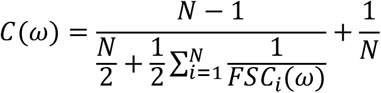

This relationship of the FSCoh with the FSC provides conceptual grounding and supports robust resolution thresholding in multi-map analysis. A full derivation and theoretical formalization are provided in the FSCoh Supplementary Material Sections 1 to 3.

### 4.3. FSCoh Thresholding

A powerful application of FSCoh is its ability to define a resolution threshold below which the information is not consistently preserved across reconstructions. The one-to-one relationship between FSCoh and FSC facilitates straightforward translation between thresholds, with the latter being the gold standard in the field.

According to [11], significant information is present in the reconstruction up to a threshold of *FSC* = 1/7. This allows the definition of an equivalent threshold for FSCoh, expressed as a function of the number of signals *N*:

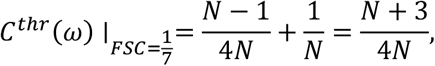

establishing that, for a given FSC threshold, the corresponding FSCoh threshold depends solely on the number of maps analysed. Furthermore, as the number of signals approaches infinity, the FSCoh threshold converges to the full-map FSC threshold, providing a direct link between the pairwise and multi-map consistency frameworks (see Table A1 in [11]). A full derivation and theoretical formalization are provided in the FSCoh Supplementary Material Section 4.

### 4.4. Statistical Maps Definition

To localize outlier densities in a target map relative to a reference pool, and to characterize any conformational or structural changes they may induce, we introduce Statistical Maps as a comparative framework.

First, every map is aligned, variance-normalized, and filtered to the resolution estimated by the FSCoh. After preprocessing, voxel-wise average, standard deviation, and median absolute deviation (MAD) maps are calculated to characterize the statistical properties of the reference pool.

The regions that significantly differ from the reference map pool in the target map are identified employing the normalized z-scores calculated as:

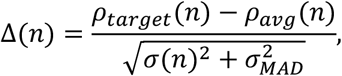

where *σ*_MAD_ is the robust estimate of the map standard deviation obtained from the MAD within the protein region. This term accounts for the specimen’s intrinsic structural variability and corrects the resulting z-scores. The protein region is defined either by a user-provided mask or by standard deviation thresholding defaulting at 1.64*σ*, which rejects the null hypothesis that a voxel originates from the noise distribution at a significance level *α* = 0.05.

To identify regions shared between the target and the reference pool, we define a coincident region as the set of voxels that are both statistically consistent with the reference variability and exhibit high contrast. Statistically consistent regions are those whose normalized z-scores lies below a specified threshold, while high-density regions are selected by applying a standard-deviation-based threshold to both average and target:

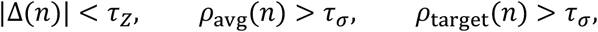

where *τ*_*Z*_ denotes the z-score threshold defining statistical consistency, and *τ*_*σ*_ is the standard-deviation-based intensity threshold applied to both the average and target maps.

### 4.5. Enhancement of Obscured Features

In some datasets, ligand density is observed with reduced contrast than the surrounding protein density, resulting in weak and poorly defined features. Such behaviour may arise from partial occupancy or other factors that reduce the contribution of the ligand signal to the reconstruction. Our objective is to recover these obscured features by removing the contribution of the reference state from the region of interest, thereby restoring a representation that better reflects the underlying ligand density.

To achieve this, we estimate the contribution of the reference state within the region of interest through a weighting factor. The weighting factor is derived by comparing voxel-intensity distributions in the previously identified coincident and different regions. To ensure a robust comparison, intensity measurements are based on a high fixed percentile (95th percentile by default), focusing the analysis on the brightest voxels within each region. Because background intensity contributes an additive component to the measured signal, percentile ratios are calculated after subtraction of the local background level. This background is estimated from a region surrounding the protein, ensuring that the resulting weighting factor reflects signal-derived differences.

This approach reflects the characteristic behaviour of cryo-EM maps, in which well-resolved regions exhibit sharp, high-density features, whereas poorly resolved regions display attenuated maxima [6]. In addition, the percentile-based metric showed greater stability with respect to variations in the shape of the detected region (see Supplementary Information).

Because the regions may differ in size, intensity comparisons are carried out using bootstrap resampling. In each bootstrap replicate, equally sized random voxel subsets are drawn from both regions and their percentile values compared. The resulting distribution of ratios provides a stable, unbiased measure of contrast between regions while effectively accounting for unequal voxel counts. Thus, the weighting factor is calculated as the median intensity ratio of this distribution.

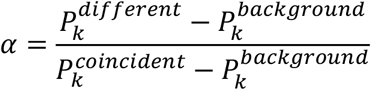

This factor is subsequently used to perform a weighted subtraction. Specifically, the average map was scaled by the estimated ratio and subtracted from the target map, attenuating the coincident regions and enhancing the obscured densities in the different region:

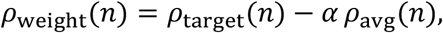

where *α* denotes the bootstrapped percentile ratio. This operation suppresses signal in the coincident regions and reveals obscured features associated with the different region. We refer to this map as the “weighted volume”, as shown in Section 2.4.

### 4.6. Code and data availability

All methods introduced in this work were implemented in the Xmipp software package [15] [16] using custom C++ modules. The source code is publicly available under an open-source license via the Xmipp public repository. Cryo-EM maps generated during and/or analysed during the current study have been deposited in the Electron Microscopy Data Bank (EMDB) under the following accession codes: EMDB-100780 (Bgal), EMDB-100781 (TRPML1 T1), EMDB-100782 (TRPML1 T2), and EMDB-100779 (PKM2 P1).

## Supporting information

Supplementary material: main

Supplementary material: FSCoh derivation

## Supplementary material

Supplementary material provides two documents. First, FSCoh Supplementary with the formal definition and derivation. Second, Supplementary Information with additional experimental results, supporting analyses, and supplementary figures that further validate and illustrate the concepts and relationships described in the main text.

## Acknowledgments

The authors acknowledge the Astex Structural Biology scientists for providing high-quality data and valuable experimental expertise, the Astex developers group for providing accessory tools for analysing and accessing the data, J. L. Vilas for discussions on the formalization of the Fourier Shell Coherence, R. S. García and M. Vinković for discussion contributions. F.P.d.I-G thanks the Sustaining Innovation Postdoctoral program from Astex Pharmaceuticals.

## Author contributions

F.P.d.I-G. conceived the method and wrote the manuscript. C.O.S.S. and J.M.C. contributed to the theoretical framework. M.S. supervised the project, wrote the manuscript and contributed to the validation of the method and the interpretation of results. All authors reviewed and approved the final manuscript.

