## Supplementary material: main for "Astex-Faro: improving detection of ligand binding in cryo-EM maps through statistical baseline analysis"

### 1. Supplementary tables

***Table 1: Data collection and refinement parameters.***

| Parameter | BGAL PETG | TRPML1 T1 | TRPML1 T2 | PKM2 P1 |
|---|---|---|---|---|
| Magnification | 120k x | 120k x | 120k x | 120k |
| Voltage (kV) | 300 | 300 | 300 | 300 |
| Electron exposure (e-/Å²) | 65.53 | 39.08 | 39.08 | 65.4 |
| Defocus range (µm) | -0.7 to -2.5 @ 0.2 steps | -0.8 to -2.0 | -1.0 to -2.0 | -0.8 to -2.2 @ 0.2 steps |
| Pixel size (Å) | 0.68 | 0.64 | 0.64 | 0.68 |
| Symmetry imposed | D2 | C4 | C4 | D2 |
| Number of movies collected | 562 | 5978 | 6381 | 572 |
| Initial particle images | 136013 | | | 267920 |
| Final particle images | 49895 | 439304 | 398353 | 142864 |
| Map resolution (Å) at FSC=0.143 | 2.2 | 2.2 | 2.1 | 2.7 |
| EMDB access codes | EMDB-100780 | EMDB-100781 | EMDB-100782 | EMDB-100779 |

***Table 2: Chemical structures of ligands used in this study***

| *Compound ID* | *Target* | *Original compound ID* | *Name* | |
|---|---|---|---|---|
| *B1* | *BGAL* | *PETG* | *PETG* | 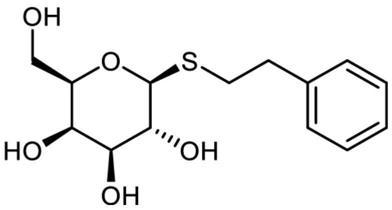 |
| *T1* | *TRPML1* | *PRU-12 /*<br>*C 13* | *N-[2-[4-(2-methoxyphenyl)piperazin-1-yl]phenyl]benzenesulfonamide* | 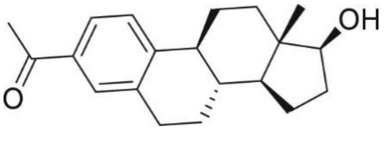 |
| *T2* | *TRPML1* | *C 10* | *N-[2-[4-(2-methoxyphenyl)piperazin-1-yl]phenyl]benzenesulfonamide* | 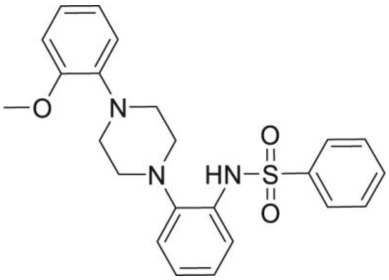 |
| *P1* | *PKM2* | *C 10* | *5-hydroxynaphthalene-1-sulfonamide* | 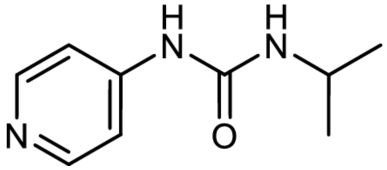 |

***Table 3: In-silico cocktailing experiment summary.***

| | | | |
|---|---|---|---|
| | | BGAL | TRPML1 |
| | | Map resolution (Å) at FSC=0.143 | |
| APO/HOLO ratio (%) | | | |
| 100 | 0 | 2.57 | 2.13 |
| 90 | 10 | 2.55 | 2.14 |
| 80 | 20 | 2.54 | 2.46 |
| 70 | 30 | 2.53 | 2.46 |
| 60 | 40 | 2.49 | 2.45 |
| 50 | 50 | 2.43 | 2.46 |
| 40 | 60 | 2.42 | 2.46 |
| 30 | 70 | 2.33 | 2.45 |
| 20 | 80 | 2.39 | 2.47 |
| 10 | 90 | 2.35 | 2.44 |
| 0 | 100 | 2.22 | 2.45 |
| | | Total number of particles | |
| | | 49895 | 40000 |

### 2. Benchmarking

#### a. Occupy

We benchmarked Astex-Faro against Occupy [1]. Occupy's graphical user interface (GUI) was used following the guidelines described in the official tutorial. Unless otherwise stated, we left all parameters at their default values or adjusted them according to the recommendations in the tutorial. For the occupancy-based weighting procedure, we evaluated several fitting modes and parameter combinations. We ultimately selected a sigmoidal fit because it consistently produced the most convincing results on visual inspection. This choice was also supported by the prior knowledge of the average ligand occupancy derived from our primary analysis, as described in the main text. We generated the final weighting using a sigmoidal fit with power = 7.4 and pivot = 0.3. Results are summarized in Figure 1.

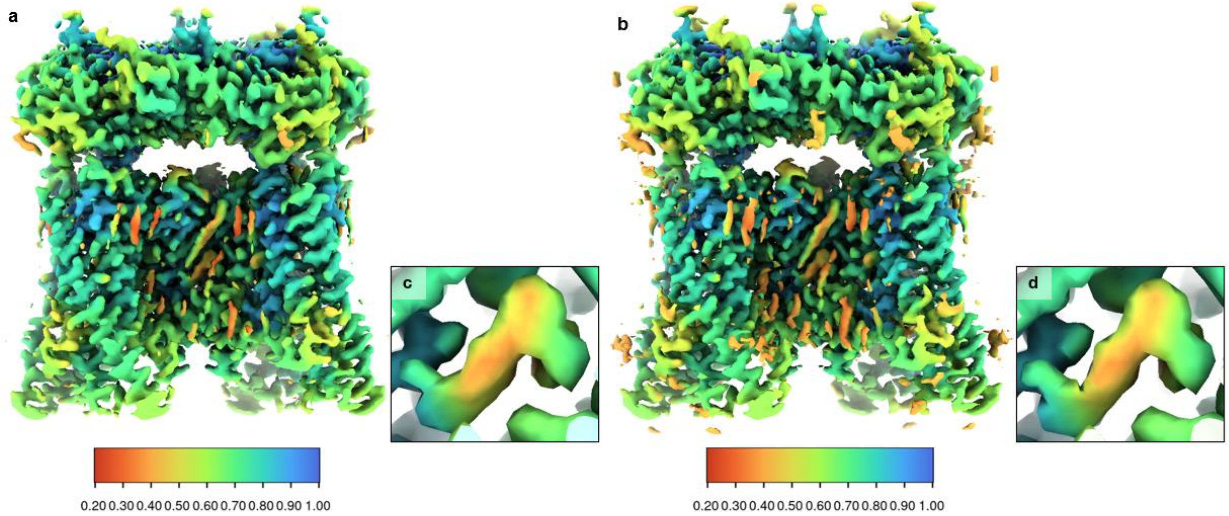

***Figure 1: Occupy-based analysis on TRPML1/compound T2 complex.*** *(a) Original reconstruction coloured according to the voxel-wise occupancy estimates generated by Occupy at threshold 0.309. (b) The same reconstruction after application of the Occupy weighting procedure using a sigmoidal fit (power = 7.4, pivot = 0.3) at threshold 0.48. (c) and (d) show a close-up view of the ligand region for every presented map respectively.*

#### b. LocScale2

We benchmarked Astex-Faro against LocScale2[2]. The program was executed following the procedure described in the official tutorial. We evaluated both available feature-enhancement map (FEM) mode together with the model-free approach. Methods requiring an atomic model were not considered, as our objective was to maintain a consistent map-only comparison with Astex-Faro, which operates exclusively on density maps. All calculations were performed using the experimental half-maps as input. No additional parameter tuning was performed. The map generated using the FEM mode were analysed with the ChimeraX scripts automatically produced by LocScale2, whereas maps generated with the model-free approach were inspected directly in ChimeraX. For visual comparison, all maps were displayed using contour levels chosen such that the highest-contrast protein regions appeared similarly represented across methods, allowing differences in contrast recovery within weaker-density regions to be assessed more easily. The FEM and model-free modes required 175 min and 196 min of computation time, respectively. Results from all evaluated modes are reported in Supplementary Material Section 2 and summarized in Figure 2.

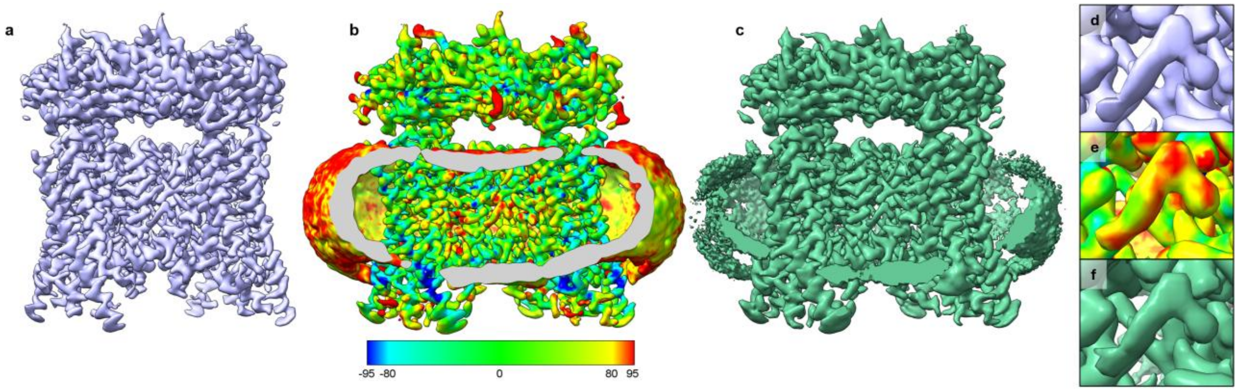

***Figure 2: LocScale2-based analysis on TRPML1/compound T2 complex.*** *(a) Original reconstruction at threshold 0.321. (b) Output map of LocScale2-FEM at threshold 0.185 coloured according to the pVDDT map. (c) Output map of Model-free LocScale2 at threshold 0.168. (d), (e) and (f) show a close-up view of the ligand region for every presented map respectively.*

#### c. Summary

Finally, we compare the performance of all methods evaluated for ligand contrast enhancement. Results are shown alongside the original reconstruction and the Astex-Faro weighted map. To assess contrast behaviour, maps are displayed using three different contour levels, selected to produce a comparable representation of the protein density surrounding the binding pocket while preserving visible differences in the ligand density. Interestingly, the LocScale2 methods also produce a smoothing effect on the protein density, affecting both ligand and protein features. This complicates the selection of contour levels for direct visual comparison. From Figure 3, Astex-Pharos can be seen to preserve ligand density details more effectively than the other methods evaluated. This is particularly evident in the ether group, where structural features remain better defined after enhancement.

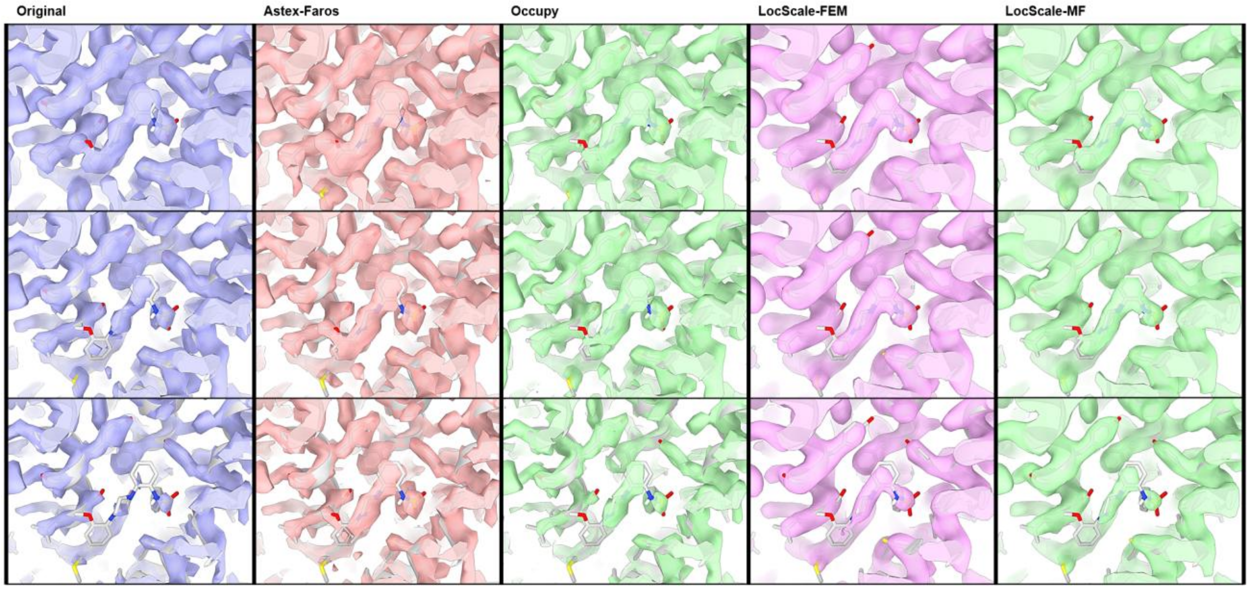

***Figure 3: Ligand close-up view on TRPML1/compound T2 complex at different thresholds.*** *Columns correspond to the original map, the Astex-Faro weighted map, Occupy, LocScale2-FEM and LocScale2-MF. Rows correspond to the three contour levels used for visual comparison. Contour levels were selected to provide a comparable representation of the protein density surrounding the binding pocket, thereby facilitating assessment of ligand contrast differences between methods.*

### 3. Analysis of weighting factor calculation stability

The calculation of the weighting factor relies on the comparison of intensities between the outlier (statistically different), coincident (statistically consistent) and background regions. These regions are defined through three different masks generated by the algorithm. However, an inherent entanglement exists in the weighting factor calculation, since the estimate depends both on the intensity differences between populations and on the extent of the masks defining the corresponding regions. As occupancy decreases, not only does the ligand signal become weaker, but the extent of the detected mask may also change. Because both effects occur simultaneously, it is difficult to determine how much of the variability in the weighting factor originates from changes in signal intensity versus changes in the detected region.

To isolate the contribution of mask estimation, we recalculated the weighting factors using a fixed reference mask obtained from a dataset composed entirely of ligand-bound particles, representing the ideal scenario for ligand-region detection. Under these conditions, the region used for the calculation remains constant across occupancies. As shown in Supplementary Figure 5, the resulting weighting factors follow a more regular trend with occupancy, indicating that part of the observed variability arises from changes in the detected masks rather than from the ligand signal alone.

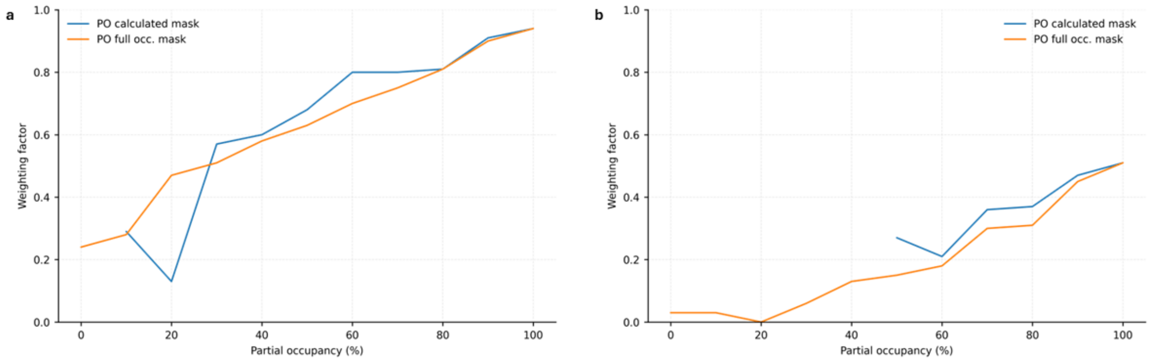

***Figure 4: Calculated weighting factors under different particle population ratios and masks.*** *Weighting factors calculated using occupancy-specific masks or a fixed reference mask obtained from a dataset composed entirely of ligand-bound particles for (a) β-galactosidase and (b) TRPML1. Values are not reported at 0% occupancy for β-galactosidase or below 50% occupancy for TRPML1 because no statistically significant regions could be reliably detected.*

### 4. Supplementary figures

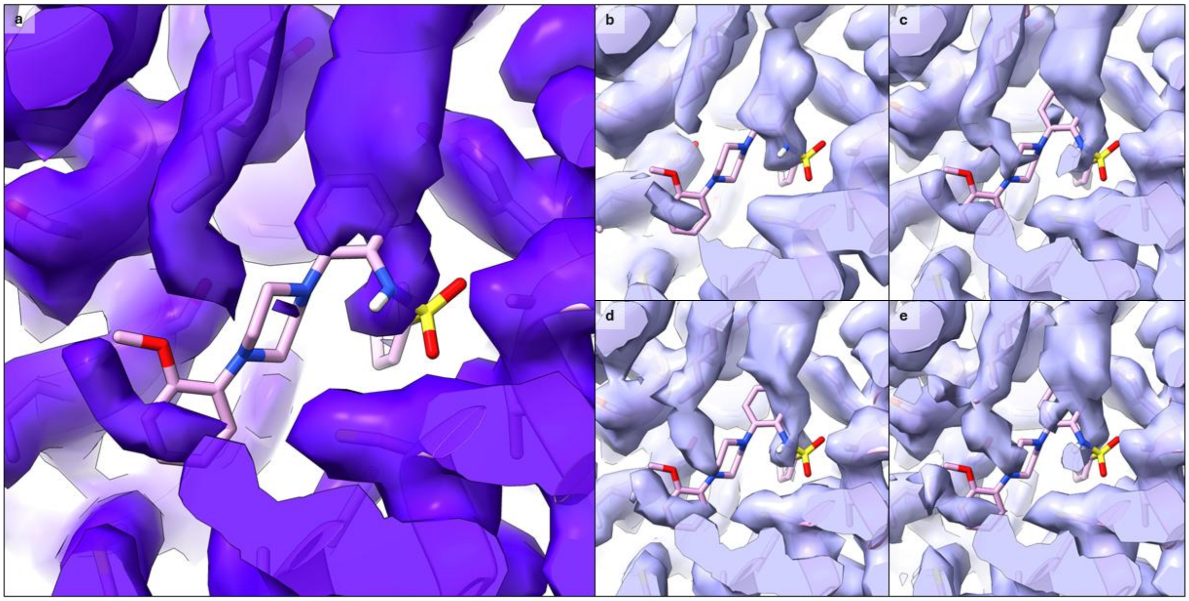

***Figure 5: Detail of the binding region in the pool of apo (unbound) maps.*** *Residual density is consistently observed within the binding pocket across the apo reference maps, which complicates the analysis by requiring the detection of subtle ligand-induced changes against a non-empty baseline. (a) Binding pocket in the average reference map calculated from the pool of apo maps. (b–e) Binding-pocket in four representative individual maps from the apo reference pool.*

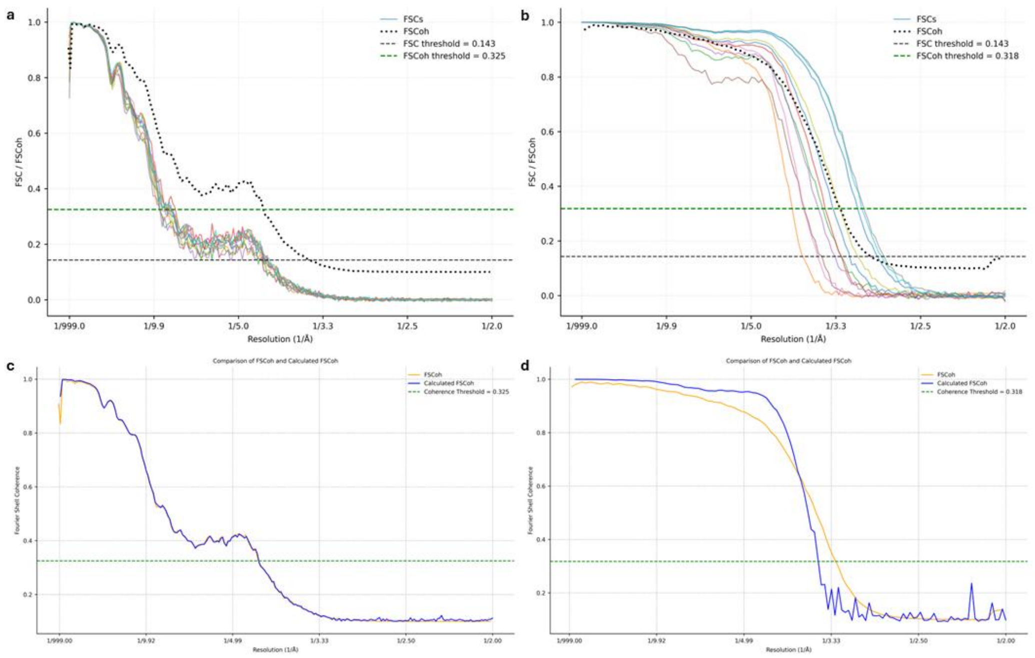

***Figure 6: Relationship between empirical and theoretical FSCoh in β-galactosidase.*** *Two scenarios were tested: (a,c) All maps originate from the same microscope acquisition and refinement, with particles split into ten independent groups, representing an ideal case in which all maps share consistent signal. (b,d) 11 maps obtained from independent acquisitions and refinements. (a,b) In addition to the empirical FSCoh the pairwise FSC curves between all pool maps are plotted. (c,d) Comparison between empirically measured FSCoh values and those calculated from the individual FSC curves. This analysis demonstrates both the mathematical relationship between FSCoh and individual FSCs (see Eq. (3) in the main text and Fourier Shell Coherence (FSCoh): supplementary material) and the expected deviation arising from structural variability and alignment inconsistencies when reconstructions come from different acquisitions. Absolute resolution values should not be considered, as particle numbers in the first scenario are reduced due to partitioning the dataset, resulting in lower overall map quality.*

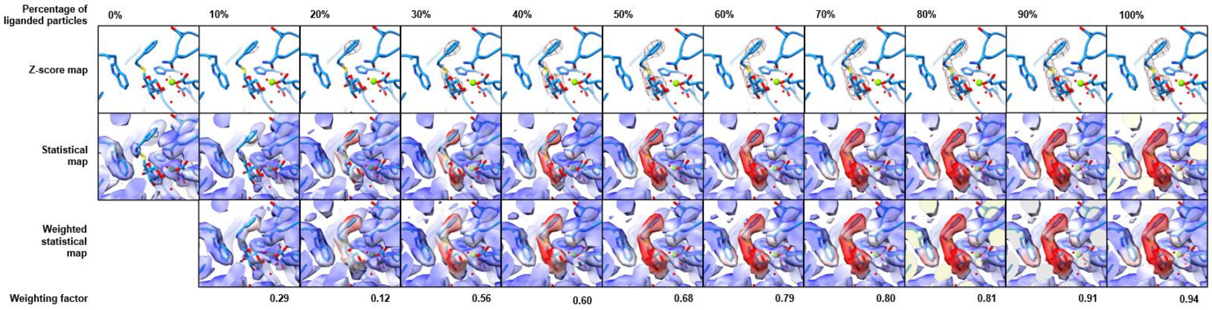

***Figure 7: Effect of partial occupancy weighting on ligand feature quality Bgal in complex with compound B1.*** *This figure is equivalent to Figure 4, showing a different orientation of the ligand. In silico mixing of apo and liganded particles, ranging from 0% to 100% liganded data, in 10% steps. Each column represents one dataset with a specific mixing ratio. First row: amount of liganded particles in each dataset. Second row: z-score map for each dataset, focused on the ligand. Third row: corresponding statistical map for each dataset. Fourth row: statistical maps weighted by the weighting factor shown in the fifth row. The weighted map for the 0% bound-particle condition is not shown, as no statistically significant regions are detected and no weighting factor can be calculated.*

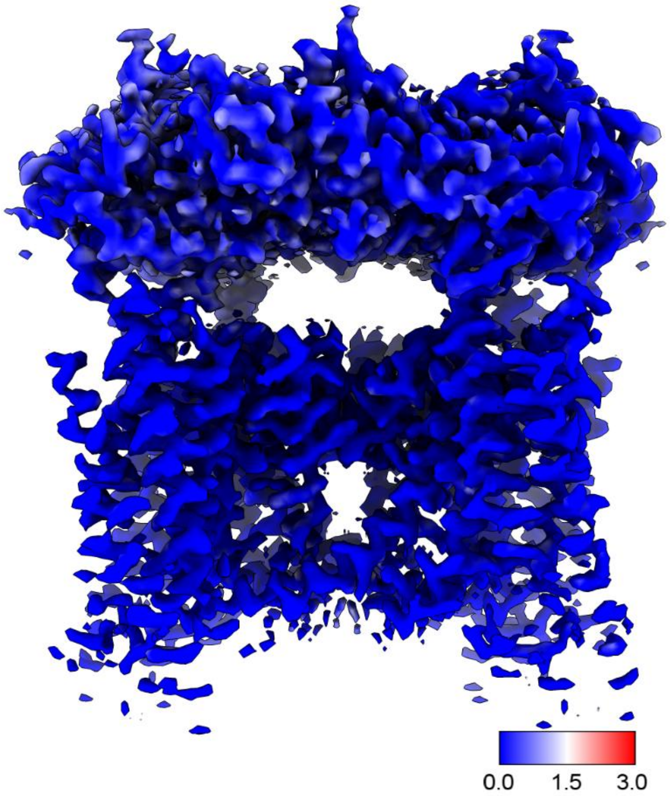

***Figure 8: Apo map compared against an apo reference pool.*** *A single apo reconstruction of TRPML1 was analysed against an apo reference pool from which it was excluded. No statistically significant features were detected, indicating that the reference ensemble accurately captures the apo state. This result further demonstrates the method's robustness to the intrinsic variability and noise present in cryo-EM reconstructions.*
