## Supplementary material: FSCoh derivation for "Astex-Faro: improving detection of ligand binding in cryo-EM maps through statistical baseline analysis"

### Fourier Shell Coherence (FSCoh): supplementary material

#### 1. Definition

To evaluate the consistency of multiple cryo-EM reconstructions, the Fourier Shell Coherence (FSCoh) is defined as a spectral metric that informs the consistency of information across a set of reconstructions. The normalized coherence, $C(\omega)$ of $N$ signals, $\{F_i(\omega)\}_{i=1}^{N}$, at frequency $\omega$ is defined as

$$C(\omega) = \frac{\int_{\Omega} |\sum_{n=1}^{N} F_n\,(\omega)|^2 d\omega}{N \int_{\Omega} \sum_{n=1}^{N} |\, F_n(\omega)|^2 d\omega} \quad (1)$$

where $F_n(\omega)$ denotes the Fourier component of the $n$-th map at frequency $\omega$ and $F_n^*(\omega)$ denotes the complex conjugate of a scalar Fourier component, averaged over shells $\Omega$ at frequency $\omega$. In the discrete case, the integral can be replaced by the dot product of two row vectors containing the Fourier components:

$$C(\omega) = \frac{|\sum_{n=1}^{N} \mathbf{F}_n^{\omega}|^2}{N \sum_{n=1}^{N} |\,\mathbf{F}_n^{\omega}|^2} \quad (2).$$

Denoting $F_j^*(\omega)$ as the complex conjugate of a scalar Fourier component and $\mathrm{F}_j^{\dagger}$ as the Hermitian conjugate for a vector $\mathrm{F}_j$ containing Fourier components over the shell $\Omega$, it is possible to rewrite Equation (3) in terms of the of dot products.

$$C(\omega) = \frac{\sum_{i=1}^{N} \sum_{j=1}^{N} \mathbf{F}_i^{\omega}\, \boldsymbol{F}_j^{\omega\,\dagger}}{\sum_{n=1}^{N} |\,\mathbf{F}_n^{\omega}|^2} \quad (3).$$

#### 2. FSCoh as SNR

Let us define the Fourier Shell Coherence (FSCoh) in terms of the SNR. To do that, we assume the observations F$i$ consists of a perfect signal $\mathrm{s}_i$corrupted by noise $\mathrm{n}_i$; that is, $\mathrm{F}_i = \mathrm{s}_i + \mathrm{n}_i$. Note that $\mathrm{s}_i$ is deterministic and $\mathrm{n}_i$ is a random variable with zero mean and standard deviation $\sigma_i$

$$C(\omega) = \frac{\sum_{i=1}^{N} \sum_{j=1}^{N} \mathbf{F}_i^{\omega}\, \boldsymbol{F}_j^{\omega\,\dagger}}{\sum_{n=1}^{N} |\,\mathbf{F}_n^{\omega}|^2} = \frac{\sum_{i=1}^{N} \sum_{j=1}^{N} (\,\mathrm{s}_i + \mathrm{n}_i)(\mathrm{s}_j + \mathrm{n}_j)^{\dagger}}{N \sum_{k=1}^{N} |\,(\mathrm{s}_k + \mathrm{n}_k)|^2} \quad (4)$$

Let us analyze the components of the previous quotient, focusing in particular on the numerator, denoted by $num$, and the denominator, denoted by $den$.

$$num = \sum_{i=1}^{N}\sum_{j=1}^{N}(\mathrm{s}_i + \mathrm{n}_i)(\mathrm{s}_j + \mathrm{n}_j)^\dagger = \sum_{i=1}^{N}\sum_{j=1}^{N}\mathrm{s}_i\,\mathrm{s}_j^\dagger + \mathrm{s}_i\mathrm{n}_j^\dagger + \mathrm{n}_i\mathrm{n}_j^\dagger + \mathrm{n}_i\mathrm{s}_j^\dagger$$
$$den = N\sum_{k=1}^{N}|\,(\mathrm{s}_k + \mathrm{n}_k)|^2 = N\sum_{k=1}^{N}|\,\mathrm{s}_k|^2 + \mathrm{n}_k\mathrm{s}_k^\dagger + \mathrm{s}_k\mathrm{n}_k^\dagger + |\mathrm{n}_k|^2 \quad (5)$$

The signals $\mathrm{s}_i$ are deterministic, whereas the noise terms $\mathrm{n}_i$ are random variables with zero mean and standard deviations $\{\sigma_i(\omega)\}_{i=1}^{N}$. We assume that the noise is isotropic and follows a symmetric distribution. Furthermore, it is plausible to consider that all signals $\mathrm{s}_i$ are identical to a reference signal $\mathrm{s}_0$, with the noise accounting for the differences among them. Also, we assume that noise terms $\mathrm{n}_i$ and $\mathrm{n}_j$ are uncorrelated for $i \neq j$. Under these assumptions, the numerator and denominator of the quotient can be approximated as

$$\mathbb{E}[num] = \sum_{i=1}^{N}\sum_{j=1}^{N}\mathbb{E}\left[\mathrm{s}_i\mathrm{s}_j^\dagger + \mathrm{n}_i\mathrm{n}_j^\dagger\right] = \sum_{i=1}^{N}\sum_{j=1}^{N}\mathbb{E}\left[|\mathrm{s}_0|^2\right] + \sum_{i=1}^{N}\mathbb{E}\left[|\mathrm{n}_i|^2\right] = N^2|\mathrm{s}_0|^2 + \sum_{i=1}^{N}\sigma_i^2$$
$$\mathbb{E}[den] = N\sum_{k=1}^{N}\mathbb{E}\left[|\mathrm{s}_k|^2 + |\mathrm{n}_k|^2\right] = N^2|\mathrm{s}_0|^2 + N\sum_{i=1}^{N}\sigma_i^2 \quad (6)$$

Equation (6) is obtained by taking the expected values of the numerator and denominator, leveraging the zero-mean property of the noise terms and their assumed uncorrelation. Thus, coherence results in

$$\mathbb{E}[C(\omega)] \approx \frac{N^2|\mathrm{s}_0|^2 + \sum_{i=1}^{N}\sigma_i^2}{N[N|\mathrm{s}_0|^2 + \sum_{i=1}^{N}\sigma_i^2]} \quad (7)$$

Finally, let us define the signal-to-noise ratio of each full map as

$$SNR_i^{full} = \frac{|s_0|^2}{\sigma_i^2} \quad (8)$$

Now it is possible to define the Coherence in terms of the SNR as

$$\mathbb{E}[C(\omega)] \approx \frac{N^2 + \sum_{i=1}^{N}\frac{1}{SNR_i^{full}}}{N\left[N + \sum_{i=1}^{N}\frac{1}{SNR_i^{full}}\right]} \quad (9)$$

#### 3. Relation between the FSCoh and the FSC

The Fourier Shell Correlation (FSC) between two signals is defined as

$$FSC(\omega) = \frac{\mathrm{F}_1\mathrm{F}_2^\dagger}{|F_1| \cdot |F_2|} (10)$$

For simplicity, it is assumed that the noise power for the two signals is approximately equal, i.e., $\sigma_1'^2 \approx \sigma_2'^2 \approx \sigma'^2$ [1]. Now, let us express the FSC in terms of the SNR, applying the same reasoning as in Equation 1

$$\mathbb{E}[FSC(\omega)] \approx \frac{\mathbb{E}[(\mathrm{s}_0 + \mathrm{n}_1)(\mathrm{s}_0 + \mathrm{n}_2)^\dagger]}{\mathbb{E}[|\mathrm{s}_0 + \mathrm{n}_1| \cdot |\mathrm{s}_0 + \mathrm{n}_2|]} = \frac{|\mathrm{s}_0|^2}{|\mathrm{s}_0|^2 + \sigma'^2} = \frac{SNR^{half}}{1 + SNR^{half}} (11)$$

From this definition, we derive the following identities.

$$SNR^{half}(\omega) = \frac{FSC}{1 - FSC} \quad \Rightarrow \quad \frac{1}{SNR^{half}} = \frac{1 - FSC}{FSC} (12)$$

$$SNR^{full}(\omega) = 2 \cdot SNR^{half} = \frac{2 \cdot FSC}{1 - FSC} \quad \Rightarrow \quad \frac{1}{SNR^{full}} = \frac{1 - FSC}{2 \cdot FSC} (13)$$

And therefore, express the FSCoh as

$$\mathbb{E}[C(\omega)] = \frac{N^2 + \frac{\sum_{i=0}^{N} 1}{SNR_i^{full}}}{N\left[N + \frac{\sum_{i=0}^{N} 1}{SNR_i^{full}}\right]} = \frac{N^2 + \sum_{i=0}^{N} \frac{1 - FSC_i}{2 \cdot FSC_i}}{N\left[N + \sum_{i=0}^{N} \frac{1 - FSC_i}{2 \cdot FSC_i}\right]} (14)$$

This establishes the relationship between the FSC and the FSCoh. Nonetheless, considering the following identity

$$N^2 = \sum_{i=1}^{N} N \frac{2 \cdot FSC_i}{2 \cdot FSC_i} (15)$$

it is possible to express the FSCoh in terms of the FSC in a more concise form

$$\mathbb{E}[C(\omega)] = \frac{\sum_{i=1}^{N} N \frac{2 \cdot FSC_i}{2 \cdot FSC_i} + \sum_{i=1}^{N} \frac{1 - FSC_i}{2 \cdot FSC_i}}{N\left[\sum_{i=1}^{N} \frac{2 \cdot FSC_i}{2 \cdot FSC_i} + \sum_{i=1}^{N} \frac{1 - FSC_i}{2 \cdot FSC_i}\right]}$$

$$= \frac{\sum_{i=1}^{N} \frac{(2N-1)FSC_i + 1}{2 \cdot FSC_i}}{N\left[\sum_{i=1}^{N} \frac{FSC_i + 1}{2 \cdot FSC_i}\right]}$$

$$= \frac{\sum_{i=1}^{N} \frac{(2N-2)FSC_i + FSC_i + 1}{2 \cdot FSC_i}}{N\left[\sum_{i=1}^{N} \frac{FSC_i + 1}{2 \cdot FSC_i}\right]} \quad (16)$$

$$= \frac{\sum_{i=1}^{N} \frac{(2N-2)FSC_i}{2 \cdot FSC_i} + \sum_{i=1}^{N} \frac{FSC_i + 1}{2 \cdot FSC_i}}{N\left[\sum_{i=1}^{N} \frac{FSC_i + 1}{2 \cdot FSC_i}\right]}$$

$$= \frac{N(N-1) + \sum_{i=1}^{N} \frac{FSC_i + 1}{2 \cdot FSC_i}}{N\left[\sum_{i=1}^{N} \frac{FSC_i + 1}{2 \cdot FSC_i}\right]}$$

$$= \frac{N-1}{\sum_{i=1}^{N} \frac{FSC_i + 1}{2 \cdot FSC_i}} + \frac{1}{N}$$

Which yields the final form of the expression

$$\mathbb{E}[C(\omega)] = \frac{N-1}{\frac{N}{2} + \frac{1}{2}\sum_{i=1}^{N} \frac{1}{FSC_i}} + \frac{1}{N} \quad (17)$$

Note that, since the coherence expected value of $N$ uncorrelated signals ($FSC(\omega) \approx 0$) is $1/N$, this expression is consistent with the expected behavior

$$\mathbb{E}[C(\omega)] \mid_{FSC_i \approx 0} = \frac{1}{N} \quad (18)$$

#### 4. Thresholding the FSCoh

According to [2], significant information is present in the reconstruction up to a threshold of $FSC = 1/7$. Following the same rationale, it is therefore possible to define an equivalent threshold for the FSCoh. To this end, we impose this condition in Equation (2):

$$C^{thr}(\omega) \mid_{FSC_i = \frac{1}{7}} = \frac{N-1}{\frac{N}{2} + \frac{1}{2}\sum_{i=1}^{N} \frac{1}{\frac{1}{7}}} + \frac{1}{N} \quad (19)$$

This leads to a definition of the FSCoh threshold as a function of the number of signals $N$

$$C^{thr}(\omega)\;|_{FSC=\frac{1}{7}}=\frac{N-1}{4N}+\frac{1}{N}=\frac{N+3}{4N}\;(20)$$

Note that this definition implies that, for an infinite number of signals, the FSCoh threshold converges to the FSC threshold for a full map (see Table A1 in [2]).

#### 5. Particularization of the FSCoh

Given the definition of FSCoh in Equation (17), there are some cases worth noting.

First, when only a single map is considered ($N = 1$), the FSCoh equals 1, since a map is inherently coherent with itself.

When two maps are compared ($N = 2$), the FSCoh is defined as:

$$C(\omega)=\frac{1}{1+\frac{1}{2}\left[\frac{1}{FSC_1}+\frac{1}{FSC_2}\right]}+\frac{1}{2}=\frac{2\cdot FSC_1\cdot FSC_2}{FSC_1+FSC_2+2\cdot FSC_1\cdot FSC_2}+\frac{1}{2}\;(21)$$

whose expected value for two uncorrelated signals ($FSC = 0$) tends to $1/2$ (which matches the expected coherence value of $1/N$).

The final case of interest occurs when the number of maps considered tends to infinity ($N \to \infty$). Let there be an infinite population of maps with an equivalent underlying signal. Thus, the population of FSC's behaves as a random variable that deviates from the mean value $\overline{FSC}$ characterized by a value $\Delta_i$ characterized by standard deviations $\{\sigma_i(\delta)\}_{i=1}^{N}$. Under this assumption, each individual FSC can be expressed as a deviation from a mean value:

$$FSC_i(\omega)=\overline{FSC}(\omega)+\Delta_i(\omega)\;(22)$$

Rewriting Equation (17) under the assumption of deviations around the mean yields

$$C(\omega)=\frac{N-1}{\frac{N}{2}+\frac{1}{2}\sum_{i=1}^{N}\frac{1}{\overline{FSC}(\omega)+\Delta_i(\omega)}}+\frac{1}{N}\;(23)$$

From now on, we will drop the frequency dependence for convenience, but it must always be understood that this dependence remains. Then, the summation in the denominator can be expressed as

$$\sum_{i=1}^{N}\frac{1}{\overline{FSC}+\Delta_i}=\frac{1}{\overline{FSC}}\sum_{i=1}^{N}\frac{1}{1+\frac{\Delta_i}{\overline{FSC}}}\;(24)$$

to be approximate using a second-order Taylor expansion

$$\frac{1}{\overline{FSC}}\sum_{i=1}^{N}\frac{1}{1+\frac{\Delta_i}{\overline{FSC}}}=\frac{1}{\overline{FSC}}\sum_{i=1}^{N}1-\frac{\Delta_i}{\overline{FSC}}+\frac{\Delta_i^2}{\overline{FSC}^2}+O(\Delta_i^3)$$
$$=\sum_{i=1}^{N}\frac{1}{\overline{FSC}}-\sum_{i=1}^{N}\frac{\Delta_i}{\overline{FSC}^2}+\sum_{i=1}^{N}\frac{\Delta_i^2}{\overline{FSC}^3}\ (25)$$

Being the deviation from the mean zero-mean independent terms and taking the expected value, the even terms cancel, leading to the following expression

$$\frac{N}{\overline{FSC}}+\frac{1}{\overline{FSC}^3}\sum_{i=1}^{N}\mathbb{E}\left(\Delta_i^2\right)=\frac{N}{\overline{FSC}}\left(1+\frac{\delta_i^2}{\overline{FSC}^2}\right)(26)$$

being $\delta_i^2$ the variance of the deviation. Finally, substituting in Equation (23):

$$\begin{aligned}\lim_{N\to\infty}C(\omega)&=\frac{N-1}{N/2+N/2\overline{FSC}\left(1+\delta_i^2/\overline{FSC}^2\right)}+\frac{1}{N}\\&=\frac{N-1}{N/2\left(1+1/\overline{FSC}\left(1+\delta_i^2/\overline{FSC}^2\right)\right)}\\&=\frac{2}{1+1/\overline{FSC}\left(1+\delta_i^2/\overline{FSC}^2\right)}\\&=\frac{2}{1+1/\overline{FSC}+\delta_i^2/\overline{FSC}^3}\end{aligned}\qquad(27)$$

Given that the fluctuations $\delta_i$ are small in absolute terms, the following simplification is applied

$$\frac{\delta_i^2}{\overline{FSC}^3}\approx 0(28)$$

leading to the reduced expression

$$C(\omega)=\frac{2\cdot\overline{FSC}}{1+\overline{FSC}}(29)$$

which matches the expression for the $FSC^{full}$ when considering infinite signals with small zero mean deviations (see Equation A4 in [2]).

#### 6. Experimental validation

To validate the preceding theoretical derivation, we designed two different experiments.

The first consists of a numerical experiment in which synthetic vectors were generated as a deterministic signal corrupted by Gaussian noise. We produced two independent noisy realizations, representing each half-map, along with their average, representing the full map. This is illustrated in Figure 1. Each vector has length $128$, with a unit signal and additive Gaussian noise of variance $\sigma^2$, serving as a discrete representation of Fourier shells. This setup mirrors the assumptions made in Equations (5)–(9), where the signal is identical across realizations and the noise is independent, isotropic, and zero-mean.

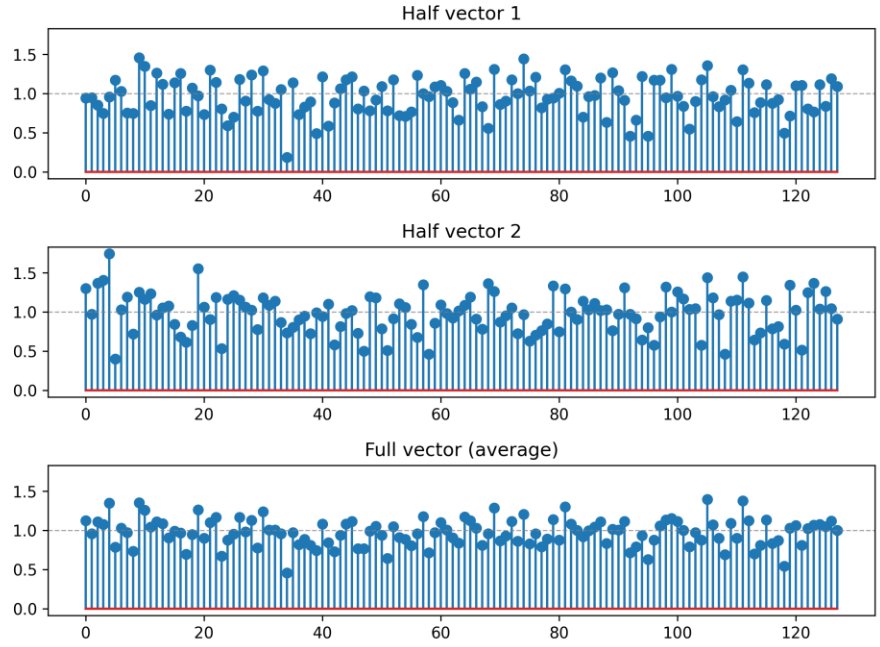


***Figure 1:*** *Example of two noisy vectors ($\boldsymbol{v_1}$ and $\boldsymbol{v_2}$) and their average $(\boldsymbol{v_1}+\boldsymbol{v_2})/\mathbf{2}$. The signal is constant and equal to 1, while Gaussian noise with standard deviation $\boldsymbol{\sigma}=\mathbf{0.25}$ is added independently to each realization.*

We generated $10^7$ vector pairs and computed several metrics, summarized in Table 1, including the mean FSC between half-maps, the empirical coherence, and the coherence estimated from individual FSCs or individual SNRs. These results closely align with the theoretical predictions based on Equations (9) and (17), confirming the derived relationship among FSCoh, FSC, and SNR.

Given that the signal and noise powers are known, the SNR for both half and full maps can be defined as in Equation (30). The empirical FSCoh ($0.96969625$) shows excellent agreement with the theoretical expectation derived from both the SNR and FSC formulations ($0.96969697$), within numerical precision. Compared with the FSCoh computed directly from the empirical FSC curves (0.969875), the agreement remains strong, though slightly less precise due to the accumulation of numerical errors across multiple operations.

This concordance reinforces the interpretation of FSCoh as a quantitative measure of consistency among multiple noisy reconstructions and demonstrates its equivalence to an SNR-based formulation.

| Metric | Value |
|---|---|
| Mean FSC | 0.94157 |
| Empirical Coherence | 0.969696 |
| Coherence from individual FSCs | 0.969875 |
| Coherence from individual SNRs | 0.969875 |
| Coherence from theoretical SNR | 0.969696 |
| Coherence from theoretical FSC | 0.969696 |

***Table 1:*** *Simulation results for* $\boldsymbol{N}_{pairs} = \mathbf{10^7}$*, vector length* $\mathbf{128}$*, and noise* $\boldsymbol{\sigma} = \mathbf{0.25}$*.*

The second experiment uses real data. Specifically, we generated ten independent reconstructions from 352,828 particles coming from a single an in-house β-galactosidase acquisition. This approach maximizes signal consistency, as all particles originate from the same microscope acquisition session, which is an ideal condition for validating the coherence-based methodology proposed in this study. The consistency across reconstructions is evident in the Fourier Shell Correlation (FSC) profiles shown in Figure 2.

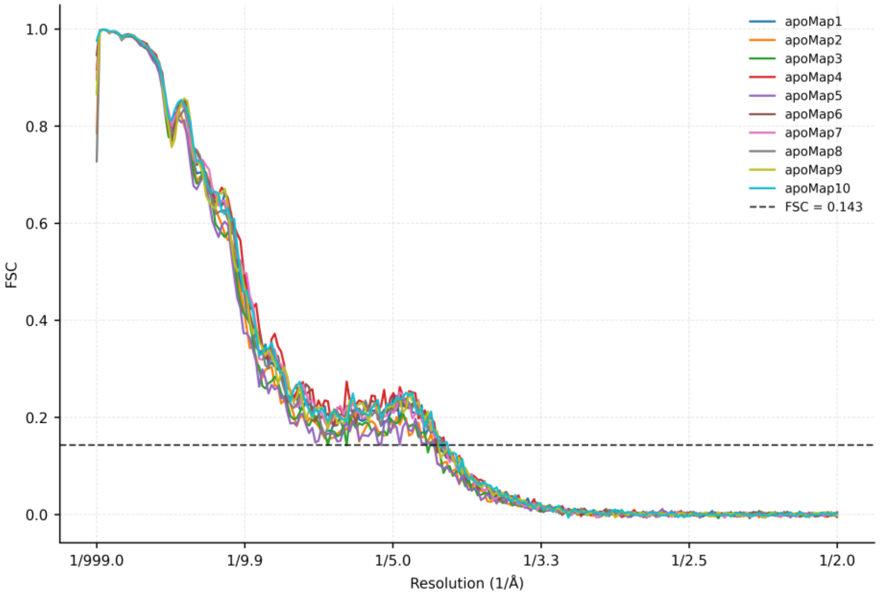


***Figure 2****: FSC profiles of the ten independent reconstructions of the Betagal dataset. The similarity among curves reflects high consistency across reconstructions.*

Consequently, we compute the empirical FSCoh by processing the set of reconstructions and derive the theoretical FSCoh from the individual FSC profiles. As illustrated in Figure 3, both curves exhibit strong agreement, thereby validating the theoretical relationship between FSCoh and FSC. Disparities at high resolution arise from numerical instabilities, particularly in regions where the FSC approaches zero.

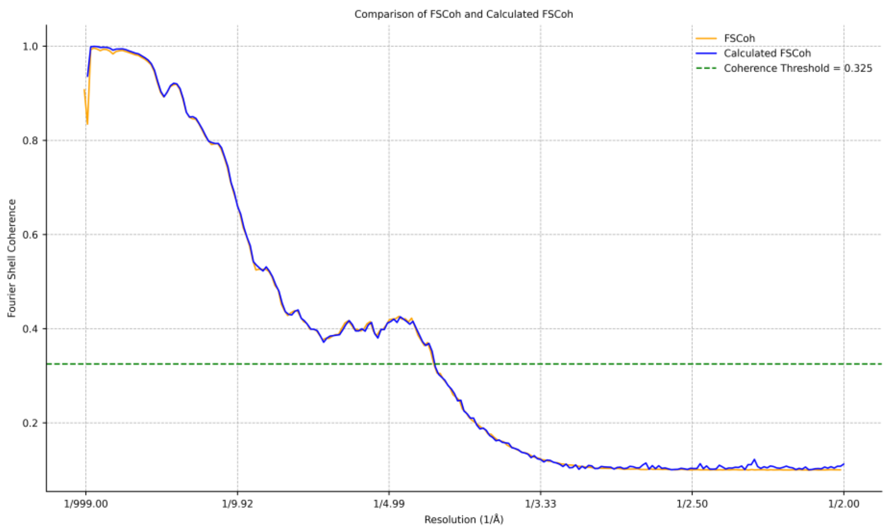


***Figure 3****: Comparison between the empirically computed FSCoh and the theoretical FSCoh derived from individual FSC profiles.*

Using real data, this experiment also validates FSCoh's ability to provide a resolution threshold. For this dataset, FSCoh indicates consistency among reconstructions up to a resolution of 4.08, consistent with the distribution of resolution values from the individual FSC curves, as shown in Figure 4.

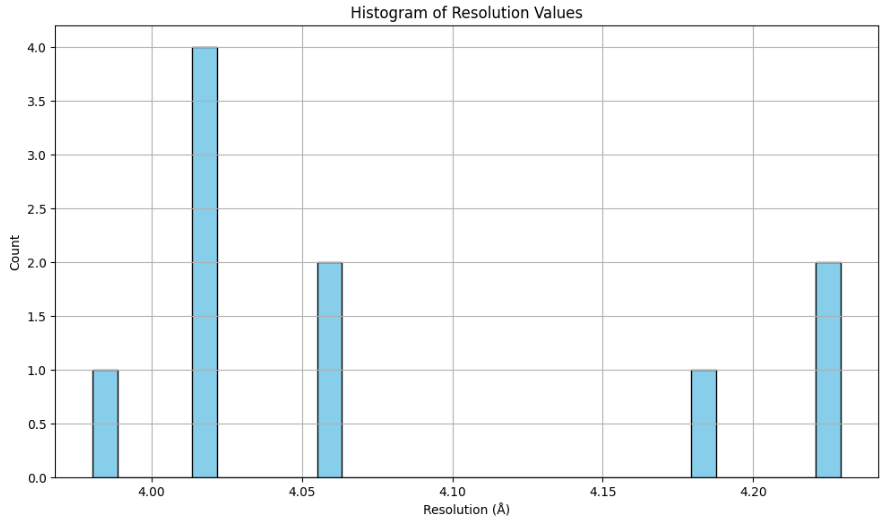


***Figure 4****: Histogram of resolution values obtained from the FSC curves of the ten reconstructions.*

These experiments thus provide a quantitative validation of FSCoh as a reliable metric for assessing the consistency and signal quality of multiple reconstructions in practical applications such as cryo-EM.
